# Mithra: An Open-Source and Cross-Platform Visualization Toolbox for Human Intracranial Recordings

**DOI:** 10.1101/2025.11.20.689539

**Authors:** Amirhossein Khalilian-Gourtani, Yasamin Esmaeili, Andrew J. Michalak, Adeen Flinker

## Abstract

Intracranial electrophysiological recordings, including electrocorticography (ECoG) and stereo-EEG (sEEG), are increasingly used across research programs to study human brain function due to their high spatiotemporal resolution. Numerous tools exist for electrode localization and visualization; however, most focus on subject-level visualization and provide only partial solutions for systematic within- and across-subject analyses. It is especially important to preserve subject-specific anatomical locations while mapping electrodes to standardized spaces for analysis across multiple subjects. To address these needs, we present **Mithra**^1^: a toolkit for visualizing intracranial recordings, implemented in both Python and MATLAB. The toolbox enables visualization of electrodes alongside the brain’s pial surface and anatomical annotations, supports localization in common average spaces such as the MNI and FreeSurfer average brains, and maintains alignment with individual anatomy. It further allows electrode projection onto the cortical surface to generate Gaussian heatmaps representing the spatial distribution of neural activity. By integrating these capabilities into a unified framework, our toolbox provides a flexible, cross-platform solution for systematic within- and across-subject electrode visualization and analysis.

## 1 Introduction

Intracranial recordings, including electrocorticography and stereo-electroencephalography, are increasingly used across clinical and research programs. Electrocorticography (ECoG) employs subdural grid and strip electrodes placed directly on the cortical surface, providing high spatial resolution over exposed regions of the brain, whereas stereo-electroencephalography (sEEG) uses depth electrodes inserted stereotactically to record from distributed cortical and subcortical structures, enabling superficial and deep sampling of neural activity. Both approaches are used in epilepsy monitoring to localize seizure onset zones and guide surgical planning. Epilepsy and cognition research programs increasingly use these recordings to study human brain function.

Accurate visualization of intracranial electrodes is essential for interpreting electrophysiological data in the context of brain anatomy. While electrode localization is typically performed in a subject’s native anatomical space, group-level analyses often require transforming electrodes into a common average space, such as the Montreal Neurological Institute (MNI) [3] or FreeSurfer’s average brain (fsaverage) [5] coordinate systems. Although linear and nonlinear transformations can approximate coordinate locations on a template brain, the a priori knowledge of the subject-specific anatomy is lost, oftentimes leading to inaccurate localization. Existing tools provide partial solutions for this process, but none integrate subject-specific anatomy within the MNI space. We provide a projection algorithm to MNI coordinate space that maintains the within-subject anatomical locations of individual electrodes, resulting in a more accurate visualization.

In recent years, there has been a growing shift toward the use of sEEG [1], driven by its ability to safely sample deep and distributed brain regions through minimally invasive implantation. However, many existing visualization tools were originally developed for ECoG and therefore provide only partial solutions for sEEG data, often lacking effective methods to visualize electrodes situated within sulci or deeper structures. To address this limitation, we provide a mapping algorithm that projects sEEG electrode coordinates onto the FreeSurfer average brain, which offers a better view of cortical folding and sulcal geometry. This enables clearer and more anatomically accurate visualization of sEEG electrodes within and across subjects.

Capturing the spatial distribution of neural activity is essential for interpreting cortical dynamics. Rather than visualizing individual electrodes, we can project activity onto the cortical surface using Gaussian heatmaps. This approach provides an intuitive visualization of the spatial distribution of neural activity recorded from discrete electrode sites. By smoothing electrode values across the cortical surface, Gaussian heatmaps produce a more continuous representation of underlying neural patterns. Despite their utility, few existing electrode visualization tools incorporate this functionality. Here, we implement Gaussian heatmaps directly on the cortical surface and provide an option to constrain the smoothing kernel to respect anatomical boundaries, ensuring that spatial interpolation remains consistent with cortical parcellations and regional organization.

## 2. Background

### 2.1 Electrode Localization Tools

Existing tools for localizing and visualizing intracranial electrodes have notable limitations. Table 1 summarizes several widely used software packages. Packages such as FieldTrip [16], NTools [20], and iELVis [8] provide MATLAB pipelines for coregistering postimplant CT or MRI to preimplant MRI and mapping electrodes onto anatomical surfaces, though they primarily focus on localization and offer limited visualization capabilities. IELU [13] and nipy [9] provide electrode localization functionality in Python and integrate with neuroimaging workflows such as MNE-Python [6]. RAVE [15] stands out for its interactive 3D visualization capabilities and its ability to combine neuroimaging with intracranial data, though it is primarily focused on localization. Meanwhile, SB-MEMA [10] introduced a framework for projecting ECoG activity onto cortical surfaces, but its lack of publicly available software limits adoption. Overall, while these methods collectively address most aspects of electrode localization with some visualization capabilities, no single package provides an integrated, open-source, and user-friendly solution for anatomically grounded visualization that is implemented across major programming environments.

**Table 1:** Overview of the existing electrode localization and visualization tools.

| Package | Platform | Notes on functionality |
| --- | --- | --- |
| FieldTrip<br>[16] | Matlab | • Postimplant CT/MRI to preimplant MRI coregistration.<br>• GUI for manual localization of electrodes in intra-operative photographs or postimplant neuroimaging. |
| IELU [13] | Python | • Postimplant CT to preimplant MRI coregistration.<br>• Automatic electrode extraction.<br>• GUI for manually tweaking electrode locations and grid information.<br>• Exporting to montage files suitable for use in MNE-python [7]. |
| NTools [20] | Matlab | • Postimplant CT to preimplant MRI coregistration.<br>• Visualizes subdural electrodes overlaid on the brain’s surface.<br>• Maps electrodes to the MNI average brain. |
| SB-MEMA<br>[10] | Matlab<br>R | • Registers ECoG data to subject-level cortical topology.<br>• Provides projection algorithm for creating surface heatmaps for ECoG recordings.<br>• No software is publicly available. |
| iELVis [8] | Matlab | • Postimplant CT/MRI to preimplant MRI coregistration.<br>• Maps ECoG electrodes to the FreeSurfer space for anatomical annotation. |
| RAVE [15] | R | • Provides display of subject cortical surface for ECoG electrode localization.<br>• Can simultaneously visualize neuroimaging and intracranial electrodes via 3D viewer. |
| nipy [9] | Python | • Postimplant CT to pre/postimplant MRI coregistration.<br>• Focused on electrode localization. |

**Table 2:** Summary of Common FreeSurfer Surface and Annotation Files.

| Category | Files and Description |
| --- | --- |
| **Pial Surfaces** | – `lh./rh.pial`: Pial surfaces (L/R)<br>– `lh./rh.sphere`: Spherical coordinate system<br>– `lh./rh.inflated`: Smoothed, inflated cortex (L/R)<br>– `lh./rh.curv`: Cortical curvature files (L/R)<br>– `lh./rh.sulc`: Sulcal depth files (L/R)<br>– `lh./rh.white`: White matter surfaces (L/R) |
| **Annotations** | – `lh./rh.aparc.annot`: Cortical labels (Desikan)<br>– `lh./rh.aparc.a2009s.annot`: Detailed atlas (Destrieux) |

### 2.2. FreeSurfer

FreeSurfer [5] is a neuroimaging software designed for analyzing and visualizing brain magnetic resonance imaging (MRI) data [5]. Many electrode localization tools use FreeSurfer for visualization and inorder to obtain anatomical annotations. We use the pial surfaces generated by FreeSurfer in our toolbox for visualization. The pial surface represents the outer boundary of the brain’s gray matter for each subject and it is extracted using the pre-implant MRI. This surface is mainly used in our visualization to show the electrode locations within the subject anatomy. Additionally, we use the spherical coordinate system provided by FreeSurfer, to transform locations between two different brains while maintaining their anatomical relevance. We use surface annotations generated by FreeSurfer to label and identify specific brain regions based on established atlases. The following list reviews the important files from the FreeSurfer ecosystem that we use in our visualization toolbox.

### 2.3. Ntools

In this manuscript, we used NTools [20] for electrode localization. NTools localizes electrodes based on post-implant CT coregistered with pre-implant MRI. Electrode positions are identified using a semi-automatic pipeline. To assign anatomical labels, NTools leverages surface annotations from FreeSurfer in the subject’s native space. NTools provides electrode coordinates in both native subject T1 space and MNI space. For mapping to MNI space, NTools estimates a volumetric transformation from the subject space to MNI space and applies it to a 3 *×*3 *×*3 voxel cube centered at each electrode, recording the mean of the transformed coordinates as the final MNI location. In our visualization tools, we use these coordinates provided by NTools, both in the subject native space and the MNI space, and extend them with additional visualization capabilities.

## 3. Methods

### 3.1. Mithra

We developed a set of visualization tools that provide a unified workflow for displaying cortical surfaces, intracranial electrodes, and anatomical annotations across both MATLAB and Python environments. The overall structure and capabilities are parallel between the two implementations. In MATLAB, visualizations are built primarily using trisurf–based rendering of cortical meshes, while the Python version leverages PyVista for rendering and surface manipulation. Mithra support loading and displaying subject-specific or template surfaces, rendering electrodes as sphere glyphs, and overlaying anatomical labels derived from FreeSurfer parcellations. Together, these visualization modules offer a flexible and consistent framework for exploring electrode placement and neural activity within the context of cortical anatomy. In the following sections, we review some of the important Algorithms and capabilities of our visualization toolbox. Open-source implementation is provided here: https://github.com/amirhkhalilian/Mithra.

### 3.2. Electrode visualization on the MNI brain

When visualizing electrodes in MNI space, a key challenge is ensuring that their projected locations accurately reflect cortical anatomy. Electrode coordinates transformed to MNI space using NTools often do not align precisely with the cortical surface, sometimes appearing displaced or floating above the brain. This issue arises because NTools relies on 3D volumetric registration to map the subject’s brain to MNI space and then applies this transformation to electrode locations. Let *T*: ℝ^3^ → ℝ^3^ be the registration transform from subject space to MNI space, **e**_*m*_ ∈ ℝ^3^ the electrode coordinate in the subject’s native space, and 𝒩 (**e**_*m*_) ⊂ ℝ^3^ the set of voxel centers in a 3 *×* 3 *×*3 cube around **e**_*m*_. Then the MNI coordinate of electrode *m* is computed in NTools as the mean of the transformed cube:

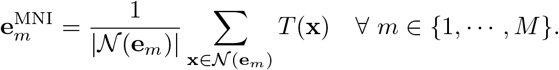

This transformation can result in electrodes being misaligned and positioned off the MNI pial surface.

A naive approach that projects electrodes to the nearest MNI pial vertex can mislocalize electrodes onto incorrect neighboring gyri, leading to distorted anatomical and functional interpretations. To mitigate this, we propose to incorporate subject-specific anatomical information to constrain electrode projection (see Algorithm 1).

Let 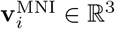 denote the coordinates of the *i*-th vertex on the MNI surface mesh.

Let 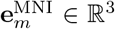 denote the coordinates of the *m*-th electrode in MNI space after the transformation from NTools, with *m* ∈ [1, · · ·, *M*] for *M* total electrodes. For each electrode, the corresponding cortical region is first identified in the subject’s native space based on anatomical labels. Here we denote by *a*_*m*_ = Atlas(**e**_*m*_) the anatomical annotation index assigned to electrode *m* in native subject space, as determined based on the per subject cortical atlas annotation. We then restrict the search for the nearest vertex on the MNI pial surface to vertices with the same cortical annotation (in Algorithm 1 the Atlas(·) function refers to the annotation of vertices in MNI space). This procedure ensures that electrodes are mapped to the closest anatomically valid location, preserving spatial fidelity while maintaining consistency with standardized MNI coordinates.

#### Algorithm 1

Electrode projection onto MNI pial surface

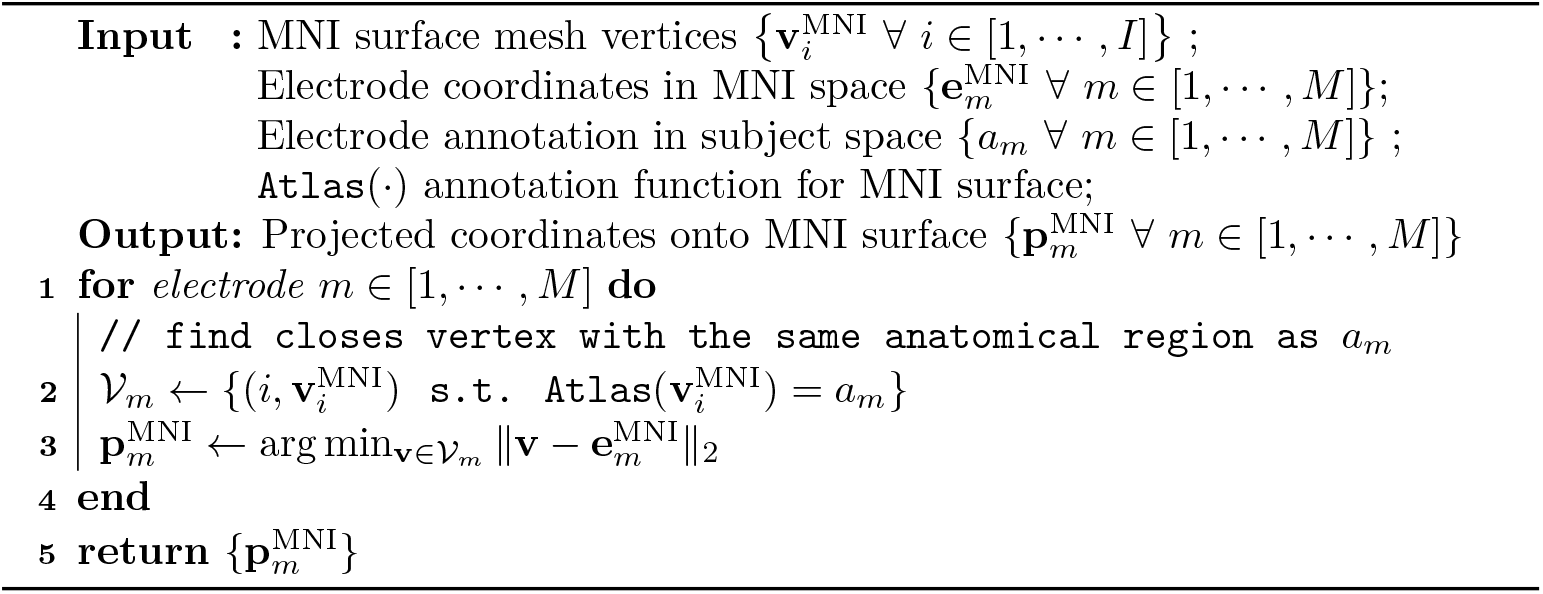

We show an example of visualizing ECoG electrodes from one subject on the MNI pial surface. The electrode coordinates on the native subject pial surface adhere to the surface and are correctly located on the corresponding gyri (Fig. 1A; color-code for each electrode follows DKT atlas). When we visualize the MNI coordinates provided by NTools on the MNI surface, we observe that the electrodes are not adhering to the surface (see Fig. 1B, frontal view) and in some cases floating in the area between gyri and mislocalized (see blue electrode at the center of zoomed region in Fig. 1B). A naive approach of moving the electrode to the closest vertex on MNI surface can lead to electrodes moving across the sulcus into adjacent gyri (Fig. 1C; blue electrode localized to pre-central gyrus in subject space moved to inferior frontal gyrus in the MNI space). By constraining the search for the closest vertex to only the ones with the same anatomical label as the electrode, the resulting electrode coordinates lie within the correct gyri surfaces (Fig. 1D; blue electrode moved to pre-central gyrus on the MNI surface).

**Fig. 1:**
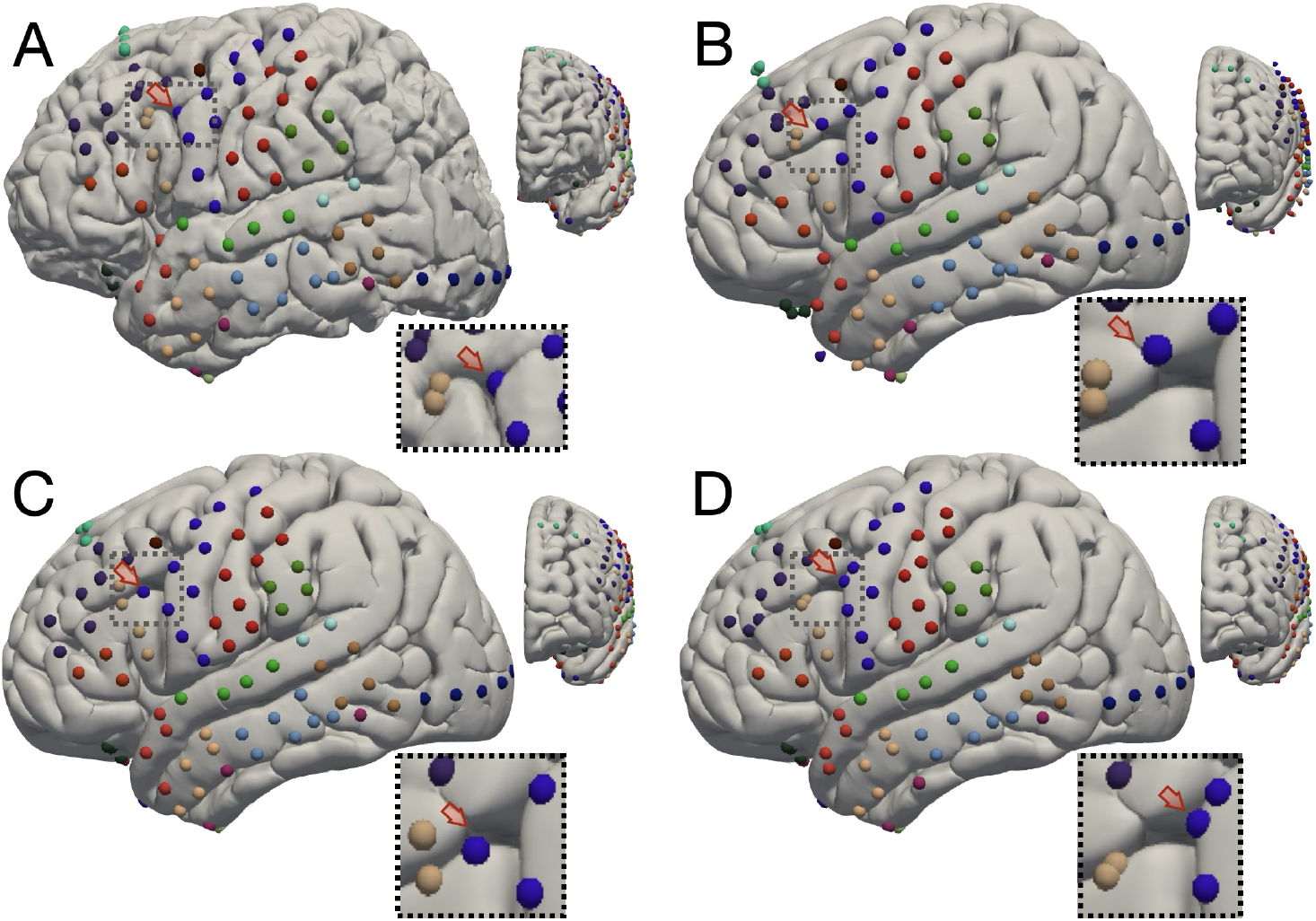
Electrode projection to the MNI surface after localization. (A) ECoG electrodes localized and visualized on the subject’s pial surface, color-coded based on their location in the DKT atlas. (B) MNI coordinates provided by NTools may not align precisely with the gyri in the MNI brain and do not necessarily adhere to the surface. (C) Naive projection directly to the closest pial vertex can result in electrodes being mislocalized to neighboring gyri. (D) Our method incorporates subject-specific anatomy to ensure electrodes are projected onto the closest vertex within the correct gyrus in the MNI space. The red arrows highlight the same electrode across all panels.

### 3.3 Plotting stereo-EEG electrodes on FreeSurfer surfaces

Mapping stereo-EEG electrodes to a common cortical surface enables consistent visualization and group-level analysis. During neurosurgical implantation, electrode trajectories are typically planned such that they sample as much cortex as possible (see Fig. 2A). This is considered a benefit of sEEG over subdural ECoG electrodes, as sEEG electrodes have the ability to not only sample gray matter at the gyral surface but also along the gyral wall and deep within the sulcus (see Fig. 2A). However, this poses a challenge to visualization and group-level analysis using the MNI pial surface because the areas inside the sulci are not visible on this surface. The FreeSurfer average brain surface (also referred to as fsaverage) provides an ideal target for such mappings, as the gyri and sulci banks are more visible on this surface.

**Fig. 2:**
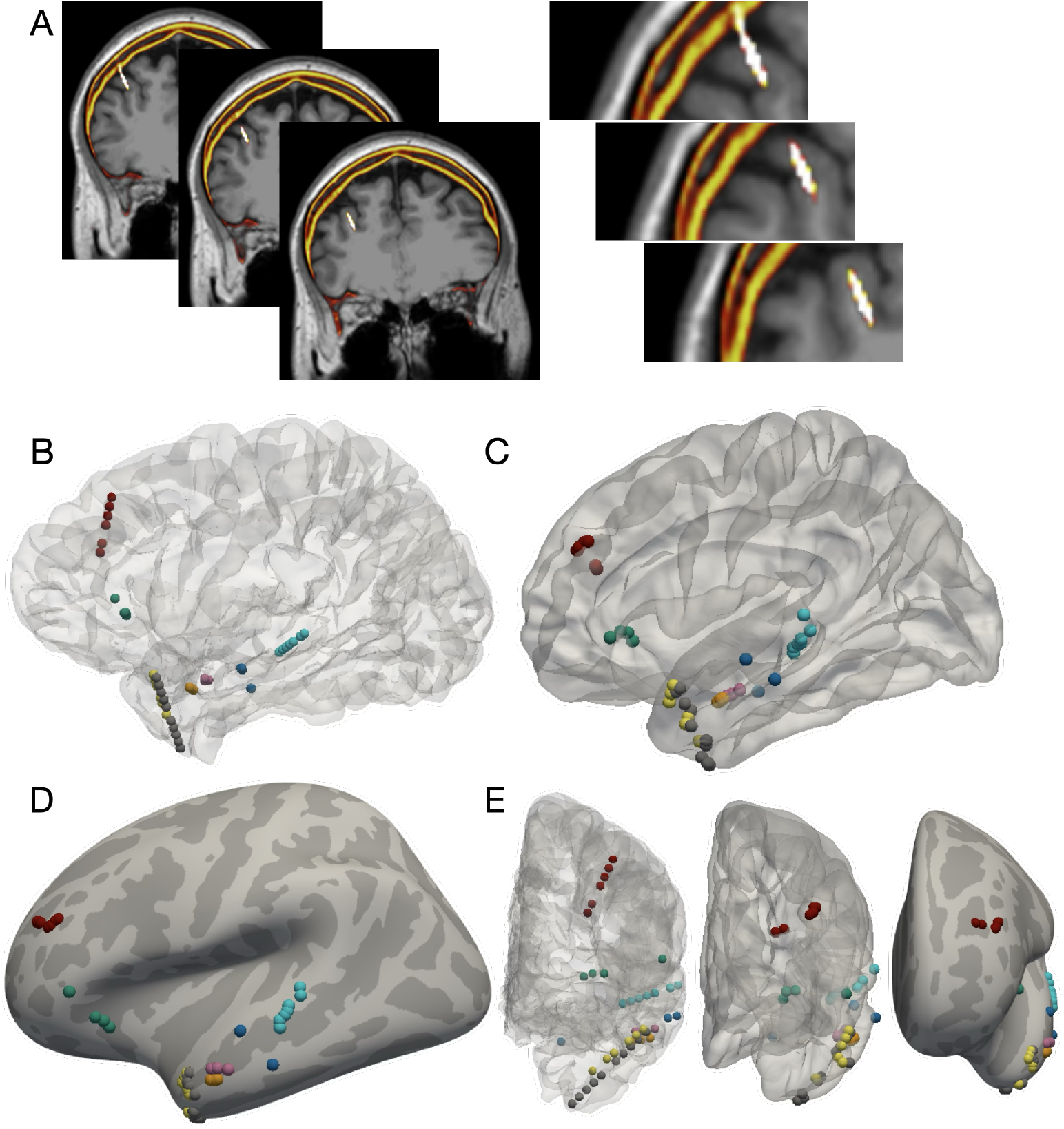
Visualization of stereo-EEG electrodes. (A) Three consecutive coronal slices of the pre-operative MRI (gray scale) and the overlaid co-registered post-operative CT (heatmap scale) show the trajectory of one sEEG lead of electrodes (the images on the right show a zoomed region around the electrodes). (B-E) Eight sets of sEEG depth electrode leads are shown on the (B) native subject cortical surface, (C) FreeSurfer average brain, and (D) FreeSurfer average inflated brain, with the corresponding frontal views depicted in (E) from left to right. From each lead we only show electrodes that are sampling cortex as spheres color-coded per lead.

To map sEEG electrodes to the fsaverage surface, we use the spherical coordinate system generated by FreeSurfer during extraction of the subject-specific pial surface. Algorithm 2 summarizes this procedure. For each electrode, the corresponding vertex on the subject’s pial surface is identified and mapped into the spherical coordinate system via index lookup. The closest point on the fsaverage sphere is then located, and its index is used to project the electrode onto the fsaverage cortical surface. This mapping preserves anatomical correspondence across subjects and supports visualization and group analyses on both the native and inflated fsaverage surfaces, as vertex indices are shared between them.

#### Algorithm 2

Mapping sEEG electrodes to Freesurfer fsaverage

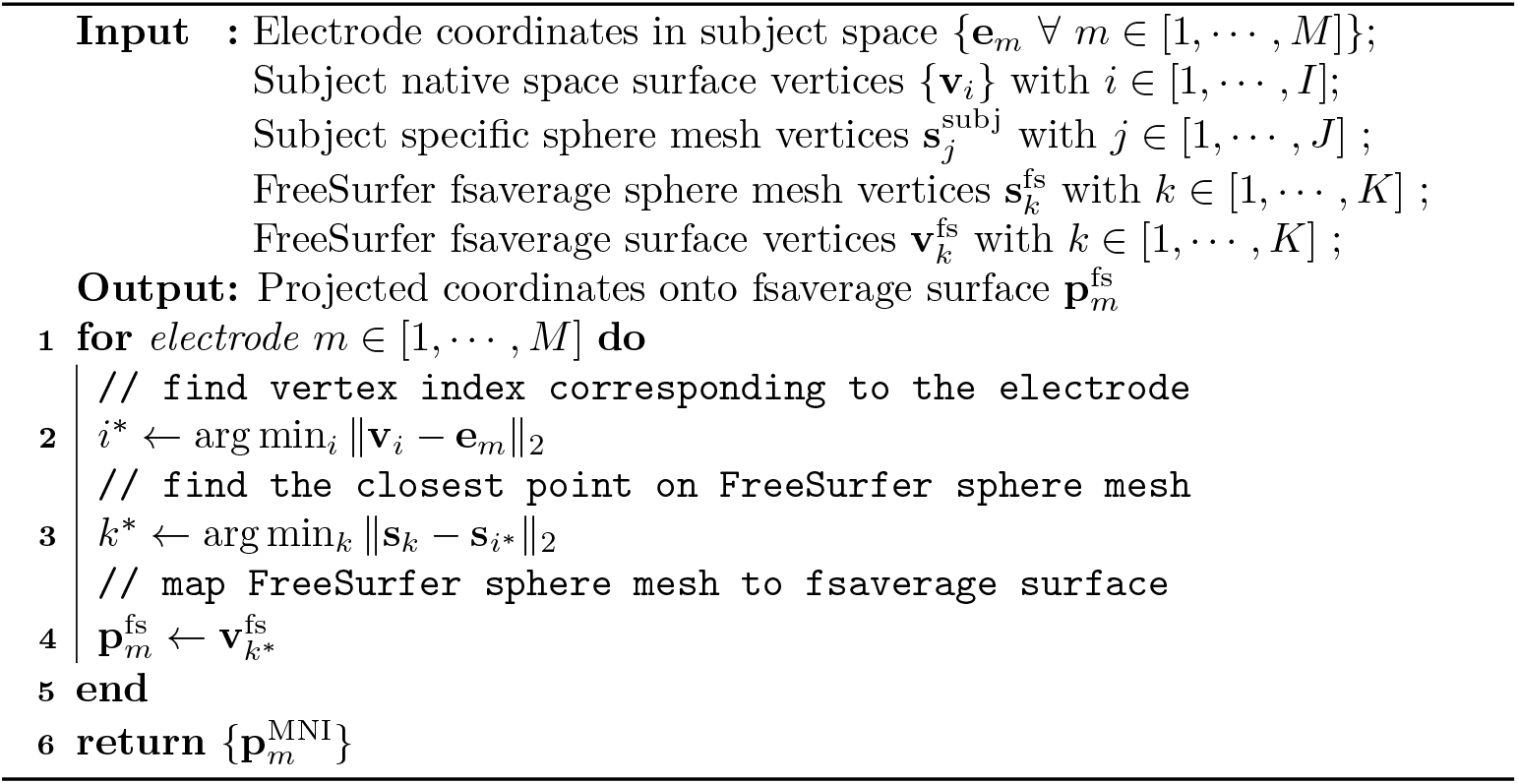

First, we show an example with three consecutive slices of the pre-operative MRI, with the co-registered CT overlaid on top (Fig.2A). These images show the trajectory of one sEEG lead and illustrate how electrode trajectories are typically planned to sample as much cortical surface as possible. Additionally, we show the other sEEG electrodes implanted in the same participant, visualized on both the native and FreeSurfer average surfaces (Fig.2B-D). Here, we display only the electrodes localized to cortical regions, excluding those in white-matter and subcortical areas. The red lead in the frontal lobes corresponds to the trajectories shown in Fig.2A.

We show visualization of the electrodes on the subject’s pial surface with increased transparency to better reveal the trajectories of different leads (Fig.2B). However, this view does not clearly show the specific gyri and sulci sampled by each electrode. By projecting the electrode locations onto the FreeSurfer average surfaces (Fig.2C,D), we can more effectively visualize their gyral assignments. Notably, on the FreeSurfer average surface, the red lead can be seen traversing along the cortical gyri (Fig.2C,E). We highlight that with the inflated surface (Fig.2D), we can visualize the electrodes on the surface with increased transparency as the gyral wall and areas within the sulcus are inflated to one smooth surface.

### 3.4. Electrode projection using Gaussian heatmaps

We provide a projection method to visualize scalar weights associated with each electrode on a cortical surface mesh. Given electrode locations and their corresponding weights, we map the weights to nearby surface vertices using a Gaussian kernel centered at each electrode’s location. We propose to constrain the projection kernel to specified anatomical regions (or any other desired atlas), using parcellation annotations. We accumulate and normalize the final projected values to account for overlapping influences across electrodes. These values are then rendered on the brain surface using a PyVista plotter or a mesh surface in Python and MATLAB, respectively. The following will describes the projection method in details (see Algorithm 3).

Let **v**_*i*_ ∈ ℝ^3^ denote the coordinates of the *i*-th vertex on a surface mesh. Let **e**_*m*_ ∈ ℝ^3^ denote the coordinates of the *m*-th electrode with associated weight *w*_*m*_, with *m* ∈ [1, · · ·, *M*], for *M* total electrodes. The distance between vertex *i* and electrode *m* is *d*_*im*_ = ∥**v**_*i*_ −**e**_*m*_∥ _2_. The Gaussian kernel projecting the influence of electrode *m* to vertex *i* is defined as

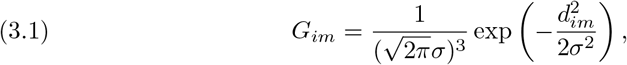

where *σ* controls the spatial spread of the kernel. The cumulative projected value at surface vertex *i* is then

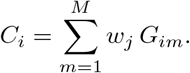

In normalized form, the color values are further normalized by the sum of kernel weights,

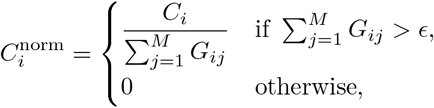

where *ϵ* is a small tolerance to avoid division by near-zero values.

Let *a*_*m*_ = Atlas(**e**_*m*_) denote the anatomical annotation index assigned to electrode *m*, as determined based on a cortical atlas. In *NTools*, for example, the anatomical region corresponding to each electrode is derived from the DKT atlas fitted to the subject-specific T1-weighted pial surface. To restrict spatial smoothing to the anatomical region associated with the electrode, we constrain the Gaussian kernel to vertices that share the same atlas annotation as the electrode. The modified kernel is defined as

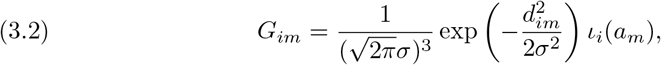

where ι_*i*_(*a*_*m*_) is an indicator function that equals 1 if vertex **v**_*i*_ belongs to the same anatomical region as *a*_*m*_, and 0 otherwise. This formulation ensures that the smoothing kernel is both spatially and anatomically constrained, thereby preventing signal leakage across cortical boundaries.

#### Algorithm 3

Project electrode weights onto mesh using Gaussian kernel.

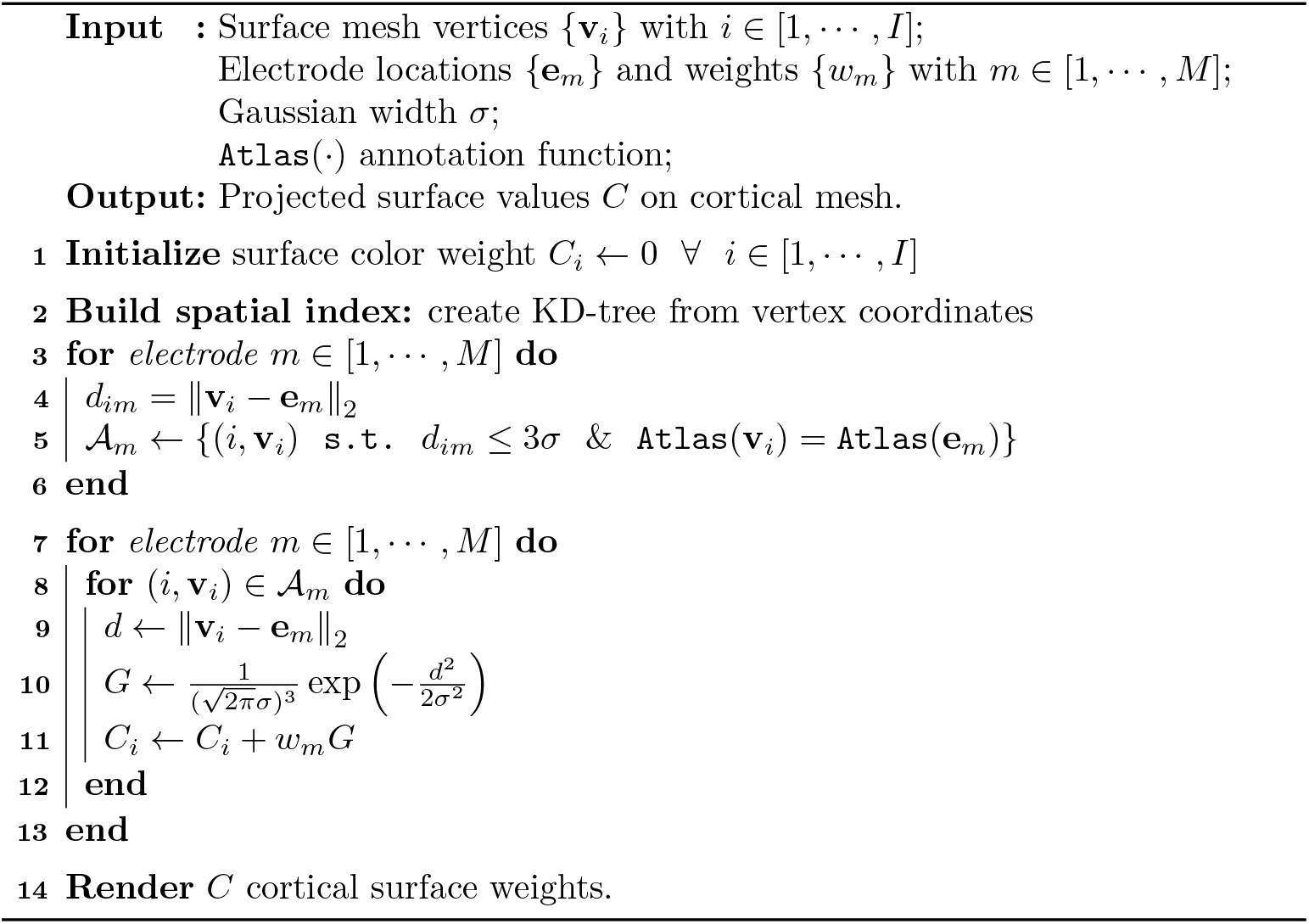

We show an example of projecting electrode weights onto the pial surface in Fig. 3. In this example we focus on the electrodes located in the middle portion of the middle superior temporal gyrus (mSTG) (Fig. 3A). Each electrode has a scalar weight associated that we aim to project onto the pial surface (Fig. 3B). We observe that an unconstrained Gaussian kernel (as in Eq. (3.1)) diffuses activity across the sulcus into adjacent gyri (Fig. 3C). In contrast, when the Gaussian kernel is constrained based on the anatomical location of each electrode (as in Eq. (3.2)), the projection with the same *σ* is correctly confined to the anatomical boundaries of the region of interest.

**Fig. 3:**
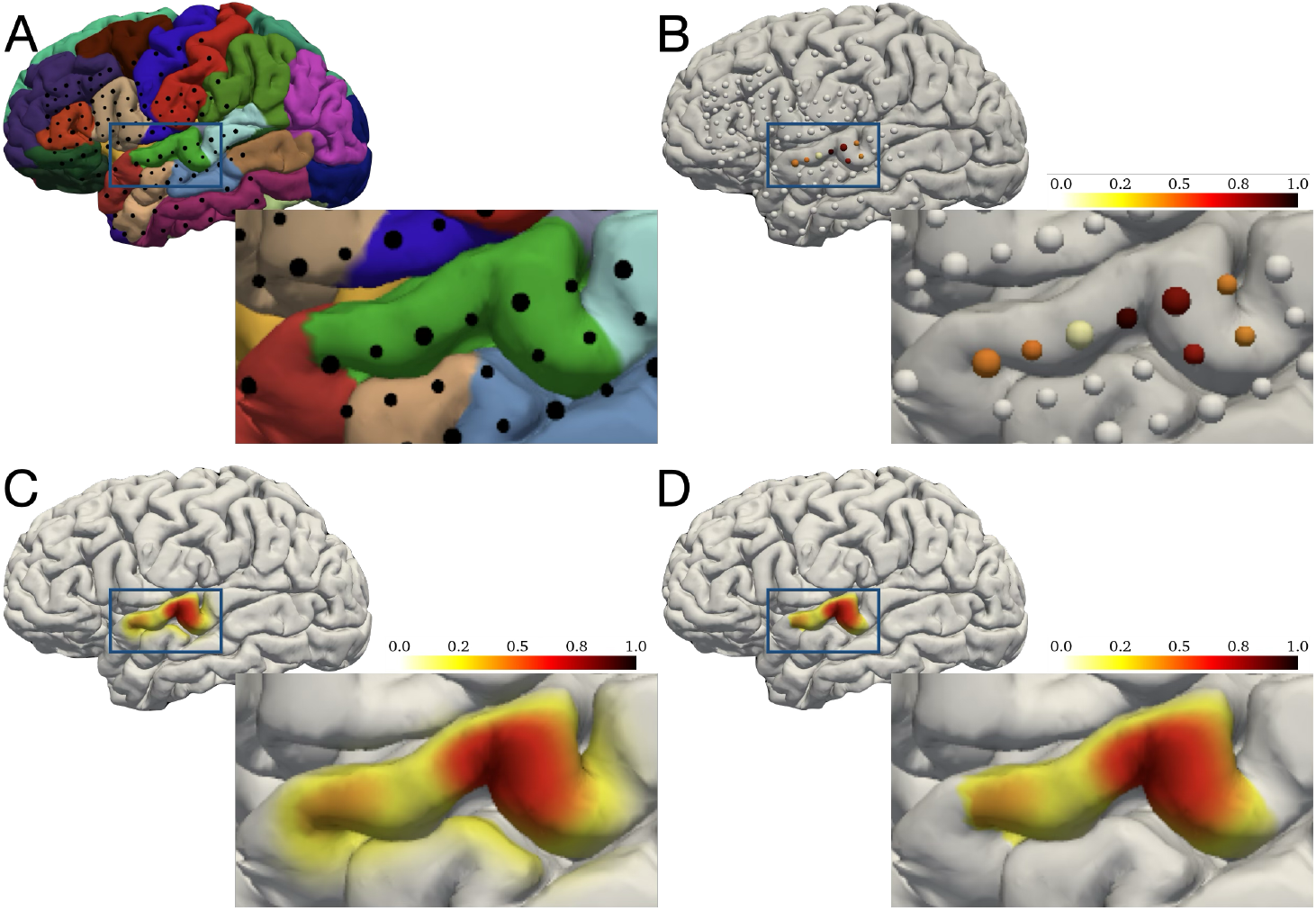
Weighted projection to the pial surface using Gaussian heatmaps. (A) A grid of electrodes is shown on the cortical surface reconstructed using FreeSurfer’s recon-all pipeline. Cortical regions are color-coded according to the Desikan–Killiany–Tourville (DKT) atlas [4], and each electrode is represented as a black sphere. (B) Example electrode weights recorded from the middle superior temporal gyrus (mSTG), with each electrode color-coded according to its weight. (C) Projection of electrode weights onto the pial surface using a Gaussian kernel unconstrained based on anatomy and with *σ* = 3 mm. (D) Projection of the same weights using a Gaussian kernel with *σ* = 3 mm, where the kernel is constrained to the anatomical region associated with each electrode.

## 4. Results

We present electrocorticographic (ECoG) recordings from a neurosurgical patient performing an auditory word repetition task [19, 11]. We analyzed high-gamma broadband activity (70–150 Hz) because of its strong correlation with underlying neuronal spiking and fMRI signals [18, 12, 14]. To characterize the spatiotemporal dynamics of auditory processing, we examined the distribution of highgamma activity relative to baseline during the listening phase of the task (Fig.4A). Neural activity was averaged within 50 ms windows centered at 100, 200, and 400 ms after stimulus onset. We observed that responses first emerged in the superior temporal gyrus (STG) around 100 ms and subsequently engaged inferior frontal regions by 400 ms. Our visualization highlights both the electrodes exhibiting task-related responses and the broader cortical regions recruited over time.

**Fig. 4:**
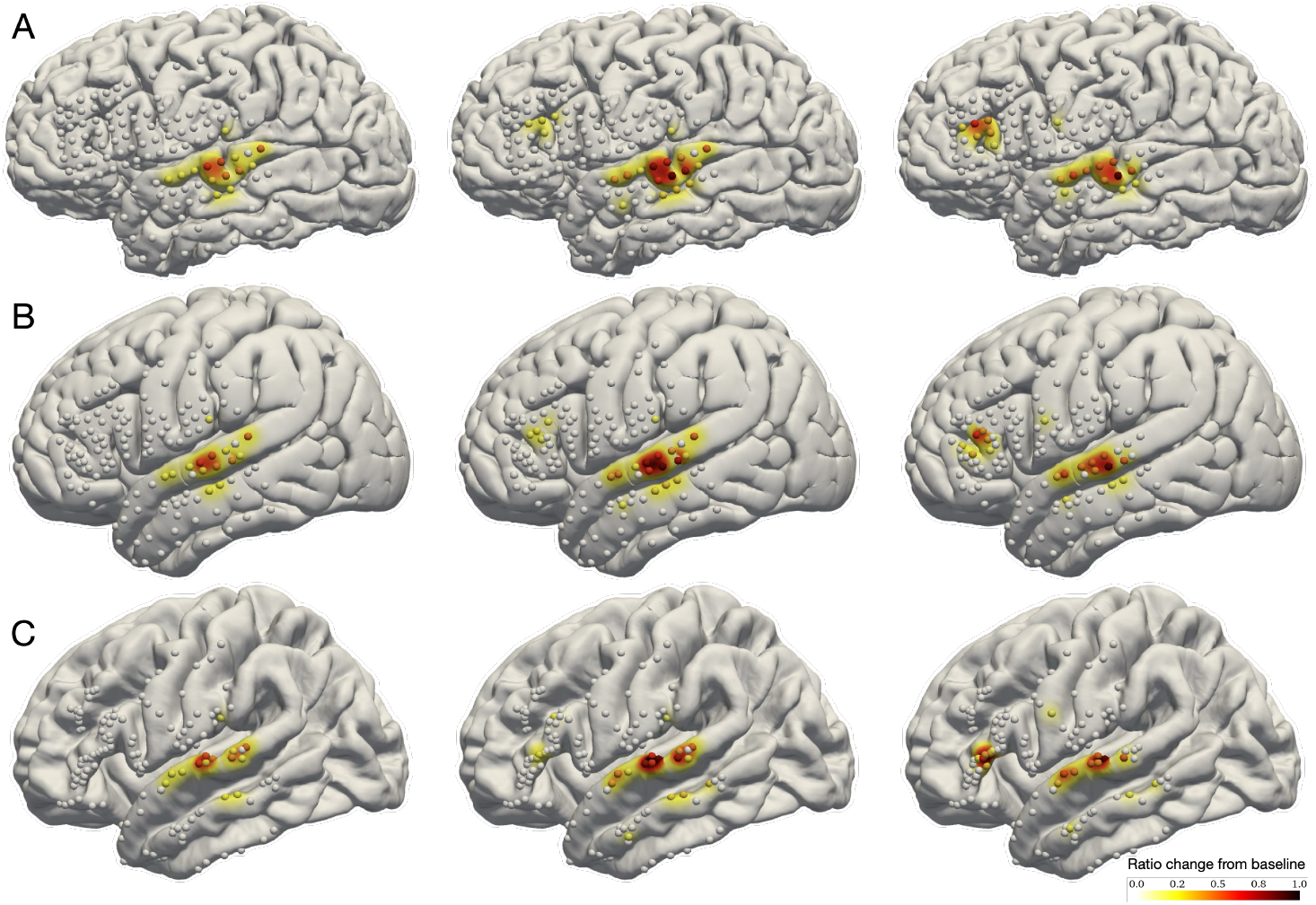
Spatiotemporal distribution of neural activity during speech perception (*N* = 1). The spatiotemporal distribution of neural activity compared to baseline in a representative participant is shown during the listening portion of an auditory word listening and repetition task. The neural activity in high-gamma (70-150 Hz) broadband, relative to baseline activity, is measured per electrode and time window. We show the activity on (A) subject native, (B) MNI, and (C) FreeSurfer average brain surfaces. Columns from left to right show 100, 200, and 400 msec after auditory stimulus onset. Electrodes are color-coded based on neural activity in units of ratio change from baseline and projected onto the respective surfaces with *σ* = 3 mm.

We then applied the transformation described in Algorithm 1 to project electrodes onto the MNI surface, ensuring accurate anatomical placement, and generated corresponding Gaussian heatmap visualizations (Fig.4B). Similarly, we used Algorithm 2 to map electrode coordinates onto the FreeSurfer average surface and applied the same Gaussian projection (Fig.4C). Across all coordinate systems, we observed consistent spatial patterns of high-gamma activity, demonstrating the robustness of our projection and visualization framework.

To demonstrate how our visualization framework generalizes across participants, we repeated the high-gamma broadband analysis for the same auditory word repetition task in five patients (previously published in [11, 17, 2]). For each participant, we projected electrode coordinates and neural activity onto both the MNI surface (Fig. 5A) and the FreeSurfer average surface (Fig. 5B) for group-level visualization of the spatiotemporal distribution. These visualizations reveal a consistent sequence of cortical engagement, with early high-gamma responses emerging in superior temporal regions and later recruitment of frontal areas. Importantly, the spatial organization of task-related activity was preserved across both coordinate systems, highlighting the robustness of our projection algorithms. This cross-participant consistency illustrates the utility of our framework for group-level intracranial electrophysiology studies.

**Fig. 5:**
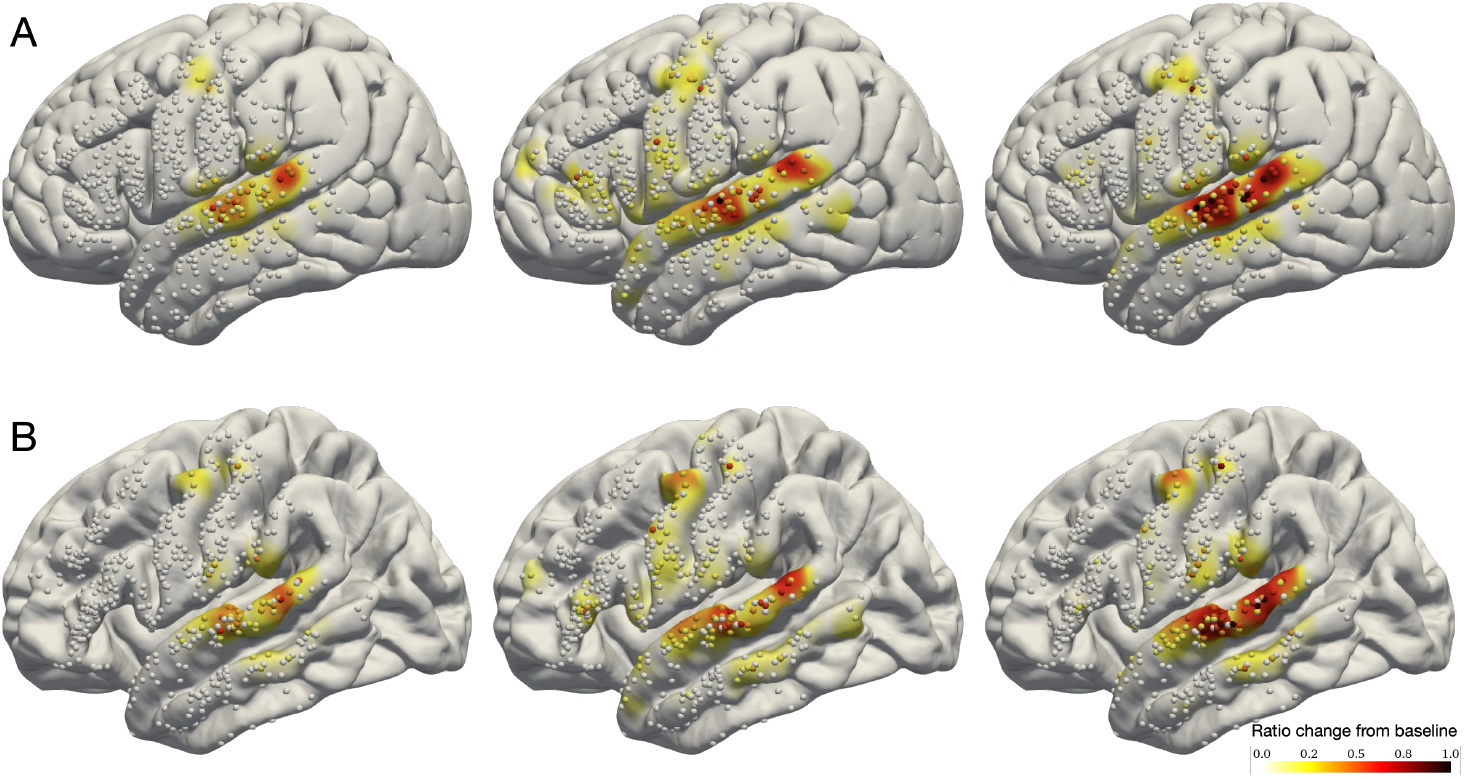
Spatiotemporal distribution of neural activity during speech perception (*N* = 5). The spatiotemporal distribution of neural activity compared to baseline across five participant is shown during the listening portion of an auditory word listening and repetition task. The neural activity in high-gamma (70-150 Hz) broadband, relative to baseline activity, is measured per electrode and time window. We show the activity on (A) MNI and (B) FreeSurfer average brain surfaces. Columns from left to right show 100, 200, and 400 msec after auditory stimulus onset. Electrodes are color-coded based on neural activity in units of ratio change from baseline and projected onto the respective surfaces with *σ* = 3 mm.

## 5. Discussion

In this paper, we present a unified framework for visualizing intracranial electrodes. Our approach addresses limitations in the existing toolkits by introducing a projection algorithm that maps electrodes to the MNI cortical surface while respecting subject-specific anatomy, a mapping algorithm for sEEG electrodes onto the FreeSurfer average brain, and a projection algorithm for creating continuous spatial representations of neural activity directly on the cortical surface via Gaussian heatmaps. Our visualization toolbox provides these functionalities in both Python and MATLAB with similar workflows. Together, these components provide an integrated solution for researchers working with both ECoG and sEEG data across different coding languages.

While existing tools provide robust solutions for discrete aspects of electrode localization, they often lack comprehensive, anatomically informed visualization capabilities, particularly for the increasingly prevalent sEEG recordings. By providing a projection that maintains anatomical fidelity in common average spaces, we enable more accurate cross-subject comparisons. Our toolbox offers a level of integrated functionality not currently available in a single, open-source package. This addresses the fragmentation in the field, where researchers often must chain together multiple specialized tools to achieve this visualization workflow.

The growing use of intracranial recordings in research and clinical practice demands visualization tools that are as sophisticated as the data they aim to represent. Our work provides a coherent framework that bridges the gap between subject-specific anatomy and group-level analysis, while accommodating the distinct visualization needs of both ECoG and sEEG methodologies. This open-source solution fills a critical niche in the neuroscience research. We anticipate that it will facilitate more accurate visualization, clearer communication, and novel insights into human brain function derived from intracranial electrophysiology.

## Acknowledgments

This work was supported by National Institutes of Health grants R01NS109367, R01NS115929, and R01DC018805 (A.F.) and National Science Foundation IIS-2309057.

## Footnotes

1 Open-source implementation is provided: https://github.com/amirhkhalilian/Mithra

